# A snapshot of the root phenotyping landscape in 2021

**DOI:** 10.1101/2022.01.28.478001

**Authors:** Benjamin M. Delory, Maria C. Hernandez-Soriano, Tomke S. Wacker, Anastazija Dimitrova, Yiyang Ding, Laura A. Greeley, Jason Liang Pin Ng, Jennifer Mesa-Marín, Limeng Xie, Congcong Zheng, Larry M. York

**Author notes:** Authors for correspondence: Benjamin M. Delory, Institute of Ecology, Leuphana University, Universitätsallee 1, 21335 Lüneburg, Germany, Larry M. York, Center for Bioenergy Innovation and Biosciences Division, Oak Ridge National Laboratory, Oak Ridge, TN, 37830, USA.

## Abstract

Root phenotyping describes methods for measuring root properties, or traits. While root phenotyping can be challenging, it is advancing quickly. In order for the field to move forward, it is essential to understand the current state and challenges of root phenotyping, as well as the pressing needs of the root biology community.

In this letter, we present and discuss the results of a survey that was created and disseminated by members of the Graduate Student and Postdoc Ambassador Program at the 11th symposium of the International Society of Root Research. This survey aimed to (1) provide an overview of the objectives, biological models and methodological approaches used in root phenotyping studies, and (2) identify the main limitations currently faced by plant scientists with regard to root phenotyping.

Our survey highlighted that (1) monocotyledonous crops dominate the root phenotyping landscape, (2) root phenotyping is mainly used to quantify morphological and architectural root traits, (3) 2D root scanning/imaging is the most widely used root phenotyping technique, (4) time-consuming tasks are an important barrier to root phenotyping, (5) there is a need for standardised, high-throughput methods to sample and phenotype roots, particularly under field conditions, and to improve our understanding of trait-function relationships.

---

Roots are crucial plant organs that underpin soil organisms through carbon and energy deposition via root exudation, while acquiring water and nutrients for plants (Freschet *et al*., 2021b). While the physical, chemical and biological functions of roots are now better understood, the ability to measure root properties and how they affect the functioning of ecosystems remains challenging to fill remaining knowledge gaps. Some of the most important knowledge gaps are which root traits to prioritise for breeding programs in agriculture, and which root traits influence carbon dynamics in the context of climate change in natural systems. Root phenotyping describes methods for measuring root properties, or traits, including root system architecture (e.g., branching angles, topology, classification by root type, etc.), morphology (e.g., specific root length, root tissue density, root hair length, etc.), mechanics (e.g., tensile strength, etc.), anatomy (e.g., root stele fraction, etc.), chemistry (e.g., root N concentration, etc.), physiology (e.g., nutrient uptake rates, root respiration, root exudation rates, etc.), and biotic interactions (e.g., root-associated microbes) (McCormack *et al*., 2017; Tracy *et al*., 2020; Freschet *et al*., 2021a). Methodological approaches for root phenotyping are diverse. They include image-based approaches using scanners, cameras and microscopes, as well as chemical abundance measurements based on infrared gas analysis, chromatography and mass spectrometry (van Dam & Bouwmeester, 2016; Atkinson *et al*., 2019; Wasson *et al*., 2020). Next-generation sequencing-based methods have also become very popular for characterising root-associated microbiota (Hannula *et al*., 2021) and quantifying species proportions in mixed root samples (Wagemaker *et al*., 2021). The diversity of approaches used in root phenotyping is illustrated in Fig. 1. While root phenotyping can be challenging, it is advancing quickly. In order for the field to move forward, it is therefore essential to understand the current state and challenges of root phenotyping, as well as the pressing needs of the root biology community.

**Figure 1.**
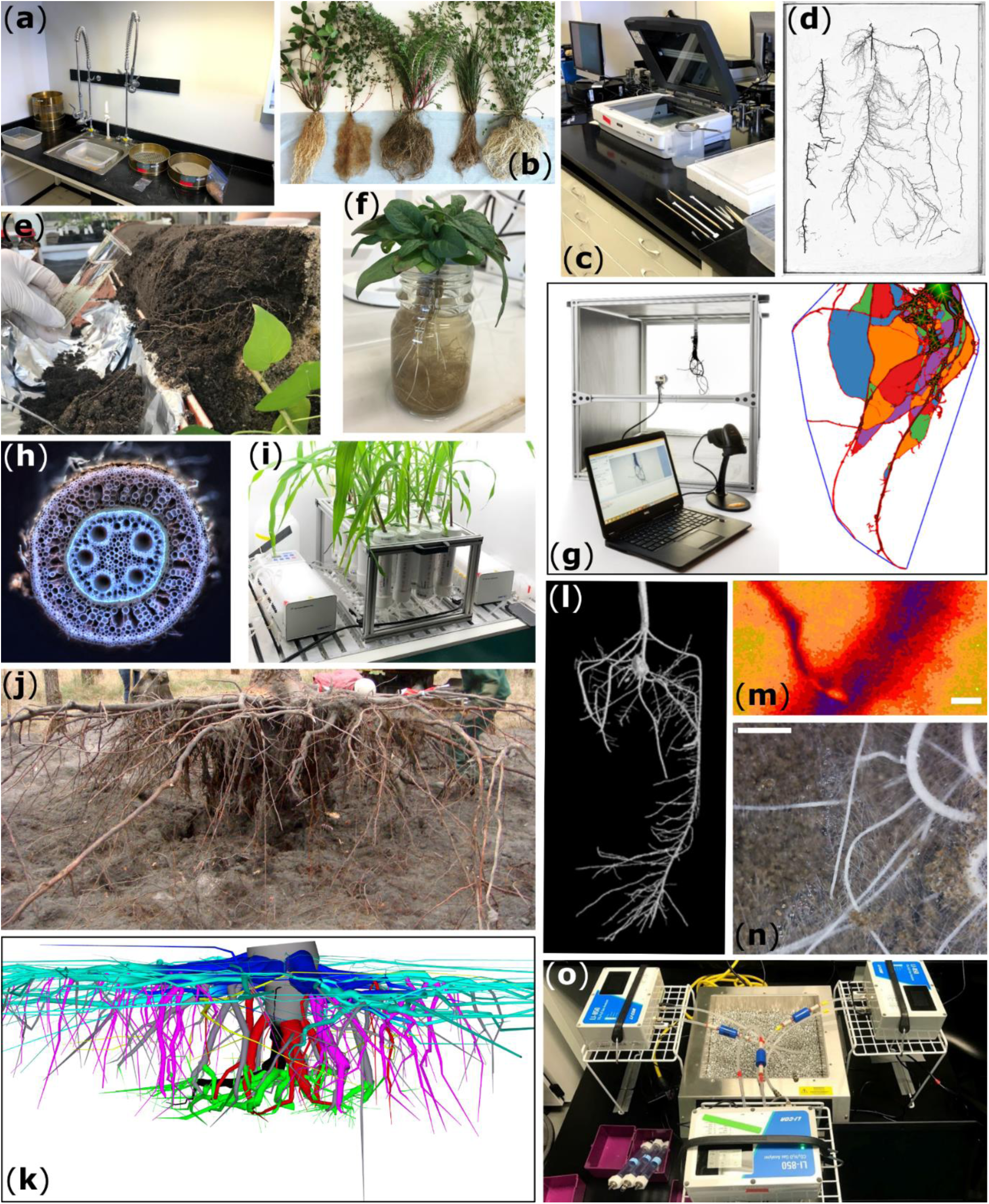
Methodological approaches for root phenotyping are diverse. (a) Root washing station (credit: Larry York). (b) Washed root systems of grassland species (from left to right: *Trifolium pratense, Galium verum, Achillea millefolium, Festuca rubra, Lotus corniculatus*) (credit: Angela Straathof). (c) Root scanning area with an Epson Expression 12000XL equipped with a transparency unit (credit: Larry York). (d) Example of an image of roots obtained with the scanner shown in panel (c). Prior to imaging, the roots were spread in a transparent plastic tray filled with a few millimetres of water (credit: Larry York). (e) Root exudate collection from tree roots (*Triadica sebifera*). A detailed description of the method can be found in (Weinhold *et al*., 2022) (credit: Sylvia Haider). (f) Collection of exudates released by *Prunella vulgaris* roots using the hydroponics-hybrid method described in (Williams *et al*., 2021, 2022) (credit: Angela Straathof). (g) Root crown phenotyping using the RhizoVision Crown platform (Seethepalli *et al*., 2020) (credit: Larry York). (h) Anatomy of a millet root imaged using laser ablation tomography (credit: Darren Wells). (i) Illustration of a high-throughput phenotyping platform (RhizoFlux) developed to measure the uptake rate of multiple ions simultaneously (Griffiths *et al*., 2021) (credit: Larry York). (j) Excavation of the root system of a 19-year-old pine tree (*Pinus pinaster*) (credit: Frédéric Danjon). (k) Digitised version of the root system shown in panel (j) (the roots have been coloured by root type). The root system was digitised using a Polhemus Fastrak low magnetic field 3D digitizer (credit: Frédéric Danjon). A description of this method as well as others to analyse the architecture of woody root systems can be found in (Danjon *et al*., 2005; Danjon & Reubens, 2008) (l) 14-day-old maize root system imaged using X-ray computed tomography (credit: https://www.nottingham.ac.uk/hiddenhalf/crop/maize.aspx). (m) In vivo imaging of pH variations in the rhizosphere of *Brachypodium distachyon* (Bd21) using planar optodes (credit: Benjamin Delory). The scale bar represents 1 mm. The image was obtained with the VisiSens A2 system from PreSens (Germany). (n) Minirhizotron image taken in a grassland field experiment with the VSI-BARTZ MS-190 camera from Vienna Scientific Instruments (Austria) (credits: Benjamin Delory, Inés Alonso-Crespo). The scale bar represents 5 mm. (o) Phenotyping platform developed to measure root respiration using a LI-COR gas analyser at a controlled temperature (credit: Larry York). A full description of this platform can be found in (Guo *et al*., 2021).

The triennial meeting of the International Society of Root Research (ISRR) was held on May 24-28, 2021 and was hosted virtually by the University of Missouri (USA). Due to the Covid-19 pandemic, a hands-on root phenotyping workshop was transitioned to a virtual “phenotyping Friday” that consisted of on-demand video tutorial collections and live discussion panels moderated by Drs. Darren Wells (University of Nottingham, UK) and Larry York (Oak Ridge National Laboratory, USA) as well as by members of the Graduate Student and Postdoc Ambassador Program. In connection with this root phenotyping workshop, the ISRR Ambassadors created and disseminated an online survey to gather key information on the current status, future directions and pressing obstacles in the field. The survey was aimed at researchers across all disciplines of plant biology. The main objectives of this survey were (1) to provide an overview of the objectives, biological models and methodological approaches used in root phenotyping studies, and (2) to identify the main limitations currently faced by plant scientists with regard to root phenotyping.

The survey was launched at the start of the ISRR11/Rooting2021 event on May 25, 2021 and closed on June 13, 2021. The web link to the questionnaire was shared by email, including ISRR members, and via the social network Twitter. A detailed description of the questionnaire and survey methodology is provided in Note S1 (Supporting information). A total of 216 researchers from 40 countries on all inhabited continents responded to the survey (Fig. 2), of which 58% attended the ISRR11/Rooting2021 event. The participants were predominantly based in the USA (21%), Germany (13%), India (11%), UK (9%) and Australia (7%) (Fig. 2). Most participants were affiliated with academia or research centres, with only 3% of participants from the industrial sector. All career stages from academia were represented, as detailed in Fig. S1.

**Figure 2.**
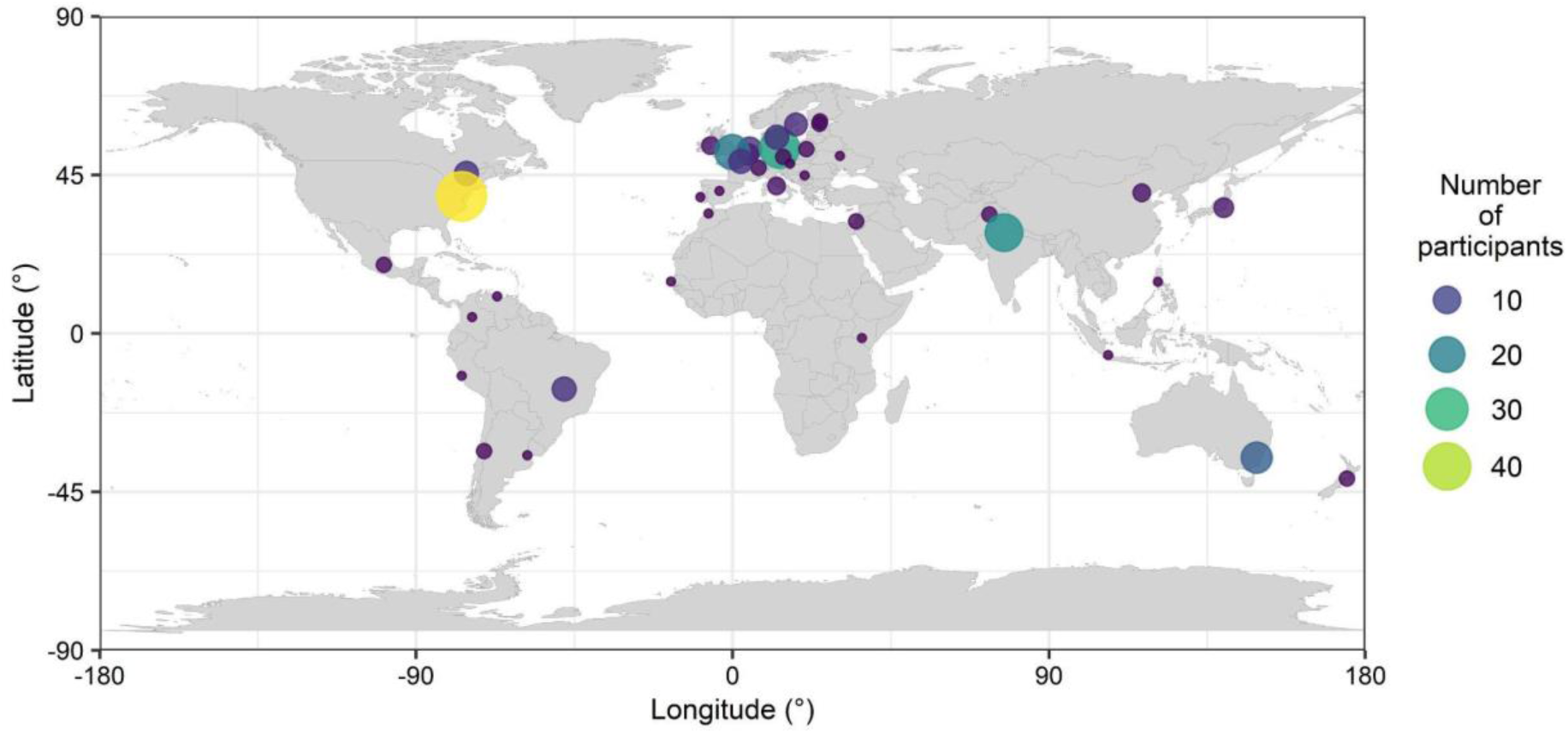
Global map depicting the distribution and number of participants who completed the online root phenotyping survey: 216 participants working in 40 countries. The circles are centred on the capital of the countries in which the participants reported working.

By providing an analysis and summary of the results of the survey, this letter aims to take a snapshot of the root phenotyping landscape in 2021. In particular, we aim to highlight the main objectives and approaches used in current root phenotyping projects, and how these will evolve to enable scientists to answer emerging questions in root research. Key highlights of our root phenotyping survey are listed in Box 1.

## Monocotyledonous crops dominate the root phenotyping landscape

The distribution of ecological habitats studied by participants was strongly skewed towards croplands (68% of the participants; Fig. 3a). This is reflected in the list of plant species most commonly used in studies using root phenotyping: maize (28%), wheat (28%), barley (16%), and rice (14%) (see Fig. S2 for a detailed description of plant species used in root phenotyping studies). Grasslands represent the second most common ecological habitat used in root phenotyping studies (17%), followed by temperate (6%), and tropical/subtropical woodlands (6%). Unsurprisingly, a significant proportion of participants (11%) use well-established lab model systems such as the model species *Arabidopsis thaliana*. Across all ecological habitats, the median soil depth to which roots are collected for phenotyping studies is 0.5 m (Fig. 3b), while a few survey participants indicated that they sampled roots in the field up to 5 m below ground.

**Figure 3.**
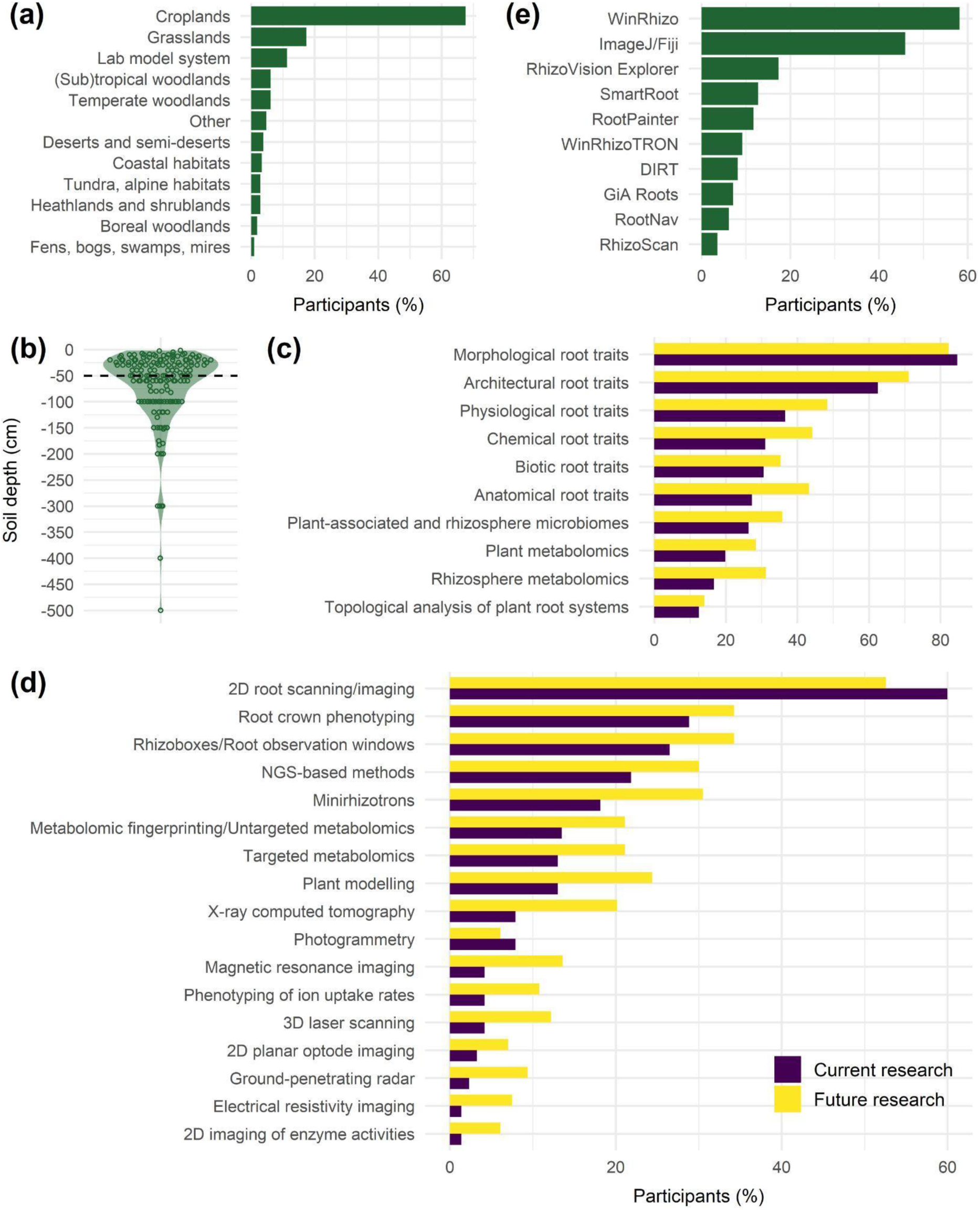
Main results of the root phenotyping survey. (a) Distribution of ecological habitats studied by survey participants in the context of root phenotyping (*n*=213). (b) Distribution of root sampling depths at which survey participants collect roots for root phenotyping (*n*=159). (c) Distribution of the main reasons why survey participants use or plan to use root phenotyping in their current and future research (current research: *n*=216; future research: *n*=215). (d) Distribution of the main root phenotyping techniques that survey participants use or plan to use in their current and future research (current research: *n*=215; future research: *n*=213). (e) Most commonly used root image analysis software tools for root phenotyping (*n*=196). In panel (a), “Lab model system” includes plants, such as *Arabidopsis*, grown under lab or greenhouse conditions (e.g., Petri dishes, hydroponics or artificial soil). The “Other” category in Panel (a) includes habitat descriptions provided by survey participants that could not be easily classified into one of the existing categories. In panel (b), the horizontal dashed line shows the median value of the distribution (0.5 m). When more than one depth value was provided by a participant, the average value was calculated and used for the analysis. Panel (e) highlights the 10 most commonly used root image analysis software tools (out of a total of 52, Fig. S6). NGS, next generation sequencing.

The survey also revealed the wide variety of growing conditions used to grow plants in root phenotyping projects, with 36% of survey participants declaring to grow plants in gel-based systems, 28% on filter papers, 79% in soil-filled pots, 40% in soil-filled rhizoboxes, 39% in hydroponics, 11% in aeroponics, 69% in growth chambers, 11% in an ecotron facility, 76% under greenhouse conditions, 36% in outdoor containers, and 63% under field conditions (Fig. S3).

## Root phenotyping is mainly used to quantify morphological and architectural root traits

Respondents reported that the main goal of using root phenotyping in their current research was to quantify morphological and architectural root traits (Fig. 3c, see also Fig. S4 for a complete description of reasons indicated by participants). We refer to McCormack *et al*. (2017) and Freschet *et al*. (2021b) for an in-depth description of the root trait categories used in our survey. Importantly, the different root trait categories were not equally represented, as physiological, chemical, biotic, and anatomical root traits seemed to be less commonly measured. Although this ranking is unlikely to change in future research (Fig. 3c), our survey revealed that a number of participants plan to expand the range of root trait categories included in their studies by also measuring physiological, chemical, biotic, and anatomical root traits. In addition to quantifying root traits, participants also highlighted another important aspect of root phenotyping, which is the characterisation of the plant and rhizosphere-associated microbiota, as well as the plant and rhizosphere metabolome (Fig. 3c). Extending root measurements to include these additional traits is important because recent research highlights the multi-dimensionality of the root economics space (Bergmann *et al*., 2020; Weigelt *et al*., 2021; Xia *et al*., 2021; Han *et al*., 2021a).

## Free, open-access and high-performance root image analysis software tools are on the rise

Among the wide variety of techniques used for root phenotyping, 2D root scanning/imaging is currently the most widely used technique (60% of participants) (Fig. 3D, see also Fig. S5 for a complete description of techniques indicated by participants). 2D root scanning/imaging is followed by root crown phenotyping (29%), which is a popular root phenotyping method in the field. In 2021, WinRhizo was still the most popular software package to analyse root images (Fig. 3e, see also Fig. S6 for a complete description of root image analysis software tools indicated by participants). However, we expect this situation to change in the future, as free, open-source, high-performance and complementary software tools such as RootPainter (Smith *et al*., 2020; Han *et al*., 2021b) and RhizoVision Explorer (Seethepalli *et al*., 2021) are becoming increasingly popular in the root research community.

Except for 2D root scanning/imaging and photogrammetry, all the other root phenotyping techniques listed in Fig. 3d will probably gain importance over time since a greater proportion of the survey participants expressed an interest in including one or more of these approaches in their future research. This is particularly true for approaches relying on root observation windows (rhizoboxes) and minirhizotrons, 3D imaging, plant modelling, or (un)targeted metabolomics. Results from our survey also highlighted the popularity of next generation sequencing-based methods in root phenotyping, which have become methods of choice to characterise the plant and soil-associated microbiota (Schöler *et al*., 2017) as well as to quantify the relative abundance of plant species in mixed root samples (Wagemaker *et al*., 2021).

## Plant modelling: an underused approach to root phenotyping?

Mathematical modelling is a powerful tool to better understand roots and rhizosphere processes and to help us identify optimal root phenotypes in contrasting environments (Roose *et al*., 2016; Postma *et al*., 2017; Pascut *et al*., 2021). Functional-structural root architecture models are particularly well suited for such endeavour (Dunbabin *et al*., 2013; Landl *et al*., 2021). However, despite the high degree of complementarity between modelling and experimental approaches, only 21% of the survey participants reported using root modelling in their research. Among those, 24% have used CPlantBox (Schnepf *et al*., 2018; Zhou *et al*., 2020), 22% OpenSimRoot (Postma *et al*., 2017), 18% RootTyp (Pagès *et al*., 2004), 13% ROOTMAP (Diggle, 1988), 13% Archisimple (Pagès *et al*., 2014), 11% GRANAR (Heymans *et al*., 2020), and 9% R-SWMS (Javaux *et al*., 2008) (Fig. S7).

## The root scientists’ wish list

When survey participants were asked about the main challenges and limitations they faced with regard to root phenotyping, 72% mentioned the fact that data collection is time-consuming (e.g., root washing). This limitation was followed by no or limited access to large and/or expensive root phenotyping equipment (e.g., X-ray computed tomography) (42%), the lack of appropriate methodologies (26%), and the lack of data and statistical analysis appropriate for root biology (21%) (Fig. S8). Between 19% and 18% of survey participants also indicated that they had no or limited access to basic root phenotyping equipment (e.g., flatbed scanner) and suffered from the lack of maintenance of available image or data analysis tools.

Hence, it is not surprising that participants emphasised the need for new methodological approaches to sample and phenotype roots, particularly under field conditions, when asked about the most urgent developments needed to enable future research plans. Overall, there was consensus on the need for standardised, high-throughput methods to collect roots, root exudates and microbial communities in the field. In addition, the need to develop advanced but affordable imaging and analysis tools for collecting data at different spatial scales was often mentioned (e.g. from root and rhizosphere metabolites to whole root systems). Non-destructive and cost-effective methods to monitor root growth and root-associated functions in systems that better represent the natural growing environment were often requested by the survey participants. The development of free, automated, accurate and easy to use image analysis software tools as well as better accessibility to root phenotyping equipment and facilities was also emphasised by survey responses. A number of participants also stressed the importance of measuring root-associated functions to enable the study of trait-function relationships in order to improve our mechanistic understanding of the roles of roots in plant and ecosystem functioning. A detailed summary of survey responses is provided in Note S2 (Supporting information).

## A way forward

This overview of the root phenotyping landscape shows a current focus of root phenotyping on the quantification of morphological and architectural traits in crops, grassland species, and the model species *Arabidopsis thaliana*. The strong dominance of monocotyledonous crops in root phenotyping studies was expected given the importance of root system traits in determining plant productivity and yield in a rapidly changing world (Tracy *et al*., 2020). The current focus on morphological and architectural traits is mainly because root phenotyping studies still rely heavily on the analysis of 2D images of roots taken with a scanner or camera, which only provide a limited set of easily measurable traits (e.g., specific root length, root tissue density, average root diameter, branching intensity, etc.) (Freschet *et al*., 2021b).

Assuming that trait data (and their associated metadata) are shared in open access global root trait databases such as FRED (Iversen *et al*., 2017) and GRooT (Guerrero-Ramírez *et al*., 2021), we see at least two benefits that would directly result from increasing the diversity of ecological habitats and the number of species included in root phenotyping studies, as well as measuring not only morphological and architectural traits, but also anatomical, mechanical, chemical, physiological and biotic traits. Firstly, it would increase the representation of species and traits that are still poorly represented in trait databases, which would allow us to better understand the mechanistic links between root traits and plant and ecosystem functioning (Freschet *et al*., 2021b). Secondly, it would improve the predictions of plant models since their parameterisation often requires information on the anatomy (e.g. stele diameter, proportion of root cortical aerenchyma, etc.) and physiological properties (e.g. root respiration, water conductivity, nutrient uptake, root exudation rate, etc.) of different root types (Postma *et al*., 2017; Schnepf *et al*., 2018; Couvreur *et al*., 2018; Heymans *et al*., 2020; Landl *et al*., 2021).

In addition to the fantastic community resources for sampling and processing roots and measuring root traits in a standardised way (Freschet *et al*., 2021a), root researchers are aided by the many technological advances in root phenotyping in recent years (Atkinson *et al*., 2019). For example, X-ray computed tomography, positron emission tomography, and magnetic resonance imaging can be used to non-destructively image roots growing in soil (Atkinson *et al*., 2019; Pflugfelder *et al*., 2021). The qualitative and quantitative analysis of root anatomy has been facilitated by the development of high-throughput root phenotyping techniques such as laser ablation tomography (Strock *et al*., 2019) or vibratome sectioning coupled with confocal microscopy (Atkinson & Wells, 2017). The development of high-resolution minirhizotrons has revolutionised our understanding of root-fungal dynamics (Defrenne *et al*., 2020), and automated root imaging platforms have enabled an in-depth characterisation of the natural variation in growth dynamics and root system architecture within species (Rellán-Álvarez *et al*., 2015; LaRue *et al*., 2021). The measurement of physiological root traits is being facilitated by the development of functional phenomics platforms for the high-throughput phenotyping of ion uptake rates and root respiration (Griffiths *et al*., 2021; Guo *et al*., 2021), as well as methods for collecting and characterising root exudates (Oburger & Jones, 2018; Uthe *et al*., 2021; Williams *et al*., 2021). Amplicon sequencing of marker genes has become a method of choice for characterising soil and plant-associated microbiota (Schöler *et al*., 2017; Nannipieri *et al*., 2019). The throughput of root image analysis pipelines has also greatly benefited from the development of tools relying on deep learning for root tip detection (Pound *et al*., 2017) or the quantification of plant-associated fungi (Evangelisti *et al*., 2021). Moreover, the complementarity that can exist between free, open access and high-performance software packages, as is the case with RootPainter (Smith *et al*., 2020) and RhizoVision Explorer (Seethepalli *et al*., 2021), holds great promise for eliminating the image analysis bottleneck that root researchers often face in their daily work. While RootPainter allows users to easily train a convolutional neural network to segment roots embedded in soil, RhizoVision Explorer allows users to automatically analyse root images providing that there is a good contrast between roots and their associated background, which is the case for the segmented images produced by RootPainter (Bauer *et al*., 2021). However, there are still some important frontiers, such as the analysis of time series for demographic studies, which require the automatic determination of the birth and death of individual roots. At this time there is more potential for more research to study root architecture and morphology than ever before in the history of plant science, from which we expect to see great gains in knowledge.

Despite these innovations, root phenotyping still poses many challenges, including the fact that it involves time-consuming tasks that may require expensive equipment and specific methodologies. Some of these challenges were highlighted by our survey participants when asked what were the most urgent developments in the field of root phenotyping that needed to be made to enable them to carry out their future research plans. In an attempt to summarise responses provided by participants, we created a root scientists’ wish list (Note S2 in Supporting information). Although not exhaustive, this list can serve as a source of inspiration for future developments and innovations to remove existing limitations, solve current challenges and enable more researchers to phenotype roots, which is essential to advance our knowledge of rhizosphere processes and species coexistence, move towards more sustainable agriculture, and develop solutions to mitigate global environmental change.

## Acknowledgements

The authors thank New Phytologist for supporting the Ambassador Program, and John Kirkegaard and Hallie Thompson for initiating the ISRR Ambassador Program in 2015. We thank Michelle Watt, Bob Sharp, Malcolm Bennett and the organisers of the ISRR11/Rooting2021 symposium from the Interdisciplinary Plant Group at the University of Missouri (Columbia, USA) and the University of Nottingham (UK). We thank Darren Wells for co-organising an excellent virtual root phenotyping workshop during the ISRR11/Rooting2021 conference with generous financial support from the International Plant Phenotyping Network. All authors would like to thank Darren Wells, Frédéric Danjon, Alex Williams, Franciska de Vries, Alexander Weinhold and Sylvia Haider for providing some of the images used in Figure 1. Finally, thanks to all those who responded to our root phenotyping survey. BMD is supported by a grant from the German Research Foundation (project 470604360). MCHS thanks Daphne Jackson Trust, BBSRC and JIC for a postdoctoral fellowship.

## Author contributions

BD analysed the data and led the writing of the manuscript, with contributions from all co-authors. All authors provided feedback before publication. The authors declare no conflict of interest.

## Data availability

Raw data and R code are available on Zenodo at https://doi.org/10.5281/zenodo.5901959.

### Box 1.

**Key highlights of our root phenotyping survey**

- Monocotyledonous crops dominate the root phenotyping landscape
- Root phenotyping is mainly used to quantify morphological and architectural root traits
- 2D root scanning/imaging is the most widely used root phenotyping technique
- Time-consuming tasks (e.g., root washing) and limited access to large and/or expensive equipment are the main barriers to root phenotyping
- Limited use of plant modelling
- Need for standardised, high-throughput methods to sample and phenotype roots, particularly under field conditions
- Need to improve our understanding of trait-function relationships

