## Supporting information for "A snapshot of the root phenotyping landscape in 2021"

**Article title**: A snapshot of the root phenotyping landscape in 2021

The following supporting information is available for this report:

**Note S1**. Survey methodology and questions asked in the root phenotyping survey.

**Fig. S1**. Detailed description of the career stages of the participants in the online root phenotyping survey.

**Fig. S2**. Detailed description of the plant species most commonly used in root phenotyping studies according to the results of the online survey.

**Fig. S3**. Range of growing conditions used to grow plants in root phenotyping studies, according to the results of the online survey.

**Fig. S4**. Distribution of the main reasons why survey participants use or plan to use root phenotyping in their current and future research.

**Fig. S5**. Distribution of the main root phenotyping techniques that survey participants use or plan to use in their current and future research.

**Fig. S6**. Root image analysis software tools used in phenotyping studies.

**Fig. S7**. Models used by survey participants in their plant modelling work.

**Fig. S8**. Limitations and challenges encountered by survey participants in their root phenotyping work.

**Note S2**. The root scientists' wish list.

**Note S1**. Survey methodology and questions asked in the root phenotyping survey.

Using a Google form, we created an online survey consisting of three sections: (1) Information about your current research (21 questions), (2) Information about your future research (5 questions), and (3) Information about the researcher (3 questions). Anyone who had the link and was interested in root research could complete the survey. A detailed description of the questionnaire is provided below.

**Section 1: Information about your current research**

1. **How often do you use root phenotyping techniques in your current research?**

   Rating question from 1 (Never) to 4 (Very often)

2. **How relevant is root phenotyping to your current research?**

   Rating question from 1 (Low relevance) to 10 (High relevance)

3. **With regard to root phenotyping, what ecological habitat(s) are you studying?**

   Multiple choice question with the following propositions:

   - Boreal woodlands
   - Coastal habitats
   - Croplands (incl. crops and pastures)
   - Deserts and semi-deserts
   - Fens, bogs, swamps, mires
   - Grasslands
   - Heathlands and shrublands
   - Temperate woodlands
   - Tropical and subtropical woodlands
   - Tundra, alpine habitats
   - Other (open field)

4. **With regard to root phenotyping, what plant species are you studying?**

   Multiple choice question with the following propositions:

   - Arabidopsis
   - Barley
   - Bean
   - Brachypodium
   - Cowpea

- Maize
- Oat
- Rice
- Sorghum
- Soybean
- Sweet potato
- Switchgrass
- Wheat
- Herbaceous species
- Shrub species
- Tree species (deciduous)
- Tree species (coniferous)
- Other (open field)

**5. Why do you use root phenotyping in your current research?**

Multiple choice question with the following propositions:

- Quantification of anatomical root traits (e.g., stele diameter, cortex fraction, etc.)
- Quantification of morphological root traits (e.g., specific root length, root diameter, root tissue density, etc.)
- Quantification of architectural root traits (e.g., branching intensity, root growth angles, deep root fraction, etc.)
- Quantification of physiological root traits (e.g., ion uptake rates, water uptake, rhizosphere acidification, etc.)
- Quantification of chemical root traits (e.g., root exudation, root N concentration, etc.)
- Quantification of biotic root traits (e.g., nodulation intensity, mycorrhizal colonisation, etc.)
- Topological analysis of plant root systems (e.g., root orders, persistent homology)
- Plant metabolomics (incl. phytohormone quantification)
- Rhizosphere metabolomics (incl. analysis of root exudates)
- Analysis of plant-associated and rhizosphere microbiomes
- I have never used root phenotyping
- Other (open field)

6. **If you use root phenotyping in your current research, under what conditions do you grow the plants?**

   Question with 11 close-ended sub-questions [Yes/No]:

   - Do you grow plants in a gel-based system (e.g., in a Petri dish or a cylinder filled with a gel medium)?
   - Do you grow plants on filter papers?
   - Do you grow plants in soil-filled pots/containers?
   - Do you grow plants in soil-filled rhizoboxes (e.g., growth container equipped with a transparent front window)?
   - Do you grow plants in hydropony?
   - Do you grow plants in aeropony?
   - Do you grow plants in a growth chamber (highly controlled environmental conditions)?
   - Do you grow plants in an ecotron (highly controlled environmental conditions)?
   - Do you grow plants in a greenhouse (semi-controlled conditions)?
   - Do you grow plants in outdoor containers (semi-controlled conditions)?
   - Do you grow plants in the field?

7. **If you use root phenotyping in your current research, what approaches or techniques do you use?**

   Multiple choice question with the following propositions:

   - Root crown phenotyping (shovelomics)
   - 2D root scanning (e.g., roots growing on Petri dishes or extracted from soil cores, pots, or monoliths)
   - Non-destructive analysis of root growth in the field using minirhizotrons
   - Non-destructive analysis of root growth using rhizoboxes or root observation windows
   - 2D planar optode imaging (pH, O2, CO2)
   - 2D imaging of enzyme activities (zymography)
   - X-ray computed tomography (3D imaging)
   - Magnetic resonance imaging (3D imaging)
   - 3D laser scanning
   - Photogrammetry
   - Ground-penetrating radar

- Electrical resistivity imaging
- High-throughput phenotyping of ion uptake rates
- Plant modelling (e.g., functional-structural plant models)
- Targeted metabolomics (i.e., quantification and identification of a small number of specific metabolites)
- Metabolomic fingerprinting and untargeted metabolomics (i.e., global and high-throughput analysis of the metabolites extracted from biological or environmental samples)
- Next generation sequencing-based methods (e.g., microbiome analysis, msGBS, etc.)
- I have never used root phenotyping
- Other (open field)

8. **If you use root phenotyping in your current research, to what soil depth do you phenotype roots?**

   Open-ended question (Please provide the maximum soil depth (in centimetres) at which you phenotype roots in the text box below. )

9. **If your root phenotyping work involves image analysis, what software tool(s) do you use?**

   Multiple choice question with the following propositions:

- ARIA
- DART
- DIRT
- DynamicRoots
- ElonSim
- EZ-Rhizo
- GiA Roots
- GLO-RIA
- GT-RootS
- GrowScreen-Root
- Growth Explorer
- IJ_Rhizo
- ImageJ/Fiji
- MorphoSnake

- NMRooting
- PlantRoot
- REST
- RhizoScan
- RNQS
- Root System Analyser
- RhizoVision Explorer
- rhizoTrak
- RootDetection
- RootGraph
- RootFly
- RootNav
- RootPainter
- RootReader2D
- RootReader3D
- RootScape
- RootSnap!
- RootTrace
- RooTrak
- RootView
- RTipC
- saRIA
- SegRoot
- Skye
- SmartRoot
- WinRhizo
- WinRhizoTRON
- Other (open field)

**10. If you use plant modelling in your current research, what model(s) do you use?**

Multiple choice question with the following propositions:

- Archisimple
- CRootBox/CPlantBox
- DigR
- GRANAR

- OpenSimRoot
- ROOTMAP
- RootTyp
- R-SWMS
- Other (open field)

**11. If you use root phenotyping in your current research, what are the limitations and/or challenges you face?**

Multiple choice question with the following propositions:

- Data collection is time-consuming
- No or limited access to basic root phenotyping equipment (e.g., 2D scanner)
- No or limited access to large and/or expensive root phenotyping equipment (e.g., 3D imaging)
- Lack of appropriate methodologies
- Lack of maintenance of available image or data analysis tools
- Lack of data and statistical analysis appropriate for root biology
- I have never used root phenotyping
- I do not face any limitations or challenges in my root phenotyping work
- Other (open field)

**Section 2: Information about your future research**

**12. What do you see as the most pressing developments in the field of root phenotyping that need to be made to enable you to carry out your future research plans?**

Open-ended question

**13. How often do you plan to use root phenotyping techniques in your future research?**

Rating question from 1 (Never) to 4 (Very often)

**14. How relevant will root phenotyping be to your future research?**

Rating question from 1 (Low relevance) to 10 (High relevance)

**15. Why do you plan to use root phenotyping in your future research?**

Multiple choice question with the following propositions:

- Quantification of anatomical root traits (e.g., stele diameter, cortex fraction, etc.)

- Quantification of morphological root traits (e.g., specific root length, root diameter, root tissue density, etc.)
- Quantification of architectural root traits (e.g., branching intensity, root growth angles, deep root fraction, etc.)
- Quantification of physiological root traits (e.g., ion uptake rates, water uptake, rhizosphere acidification, etc.)
- Quantification of chemical root traits (e.g., root exudation, root N concentration, etc.)
- Quantification of biotic root traits (e.g., nodulation intensity, mycorrhizal colonisation, etc.)
- Topological analysis of plant root systems (e.g., root orders, persistent homology)
- Plant metabolomics (incl. phytohormone quantification)
- Rhizosphere metabolomics (incl. analysis of root exudates)
- Analysis of plant-associated and rhizosphere microbiomes
- I do not plan to use root phenotyping techniques in my future research
- Other (open field)

**16. If you plan to use root phenotyping in your future research, what approaches or techniques will you use?**

Multiple choice question with the following propositions:

- Root crown phenotyping (shovelomics)
- 2D root scanning (e.g., roots growing on Petri dishes or extracted from soil cores, pots, or monoliths)
- Non-destructive analysis of root growth in the field using minirhizotrons
- Non-destructive analysis of root growth using rhizoboxes or root observation windows
- 2D planar optode imaging (pH, O2, CO2)
- 2D imaging of enzyme activities (zymography)
- X-ray computed tomography (3D imaging)
- Magnetic resonance imaging (3D imaging)
- 3D laser scanning
- Photogrammetry
- Ground-penetrating radar
- Electrical resistivity imaging

- High-throughput phenotyping of ion uptake rates
- Plant modelling (e.g., functional-structural plant models)
- Targeted metabolomics (i.e., quantification and identification of a small number of specific metabolites)
- Metabolomic fingerprinting and untargeted metabolomics (i.e., global and high-throughput analysis of the metabolites extracted from biological or environmental samples)
- Next generation sequencing-based methods (e.g., microbiome analysis, msGBS, etc.)
- I do not plan to use root phenotyping in my future research
- Other (open field)

**Section 3: Information about the researcher**

**17. In which country do you work?**

Open-ended question

**18. What is your career stage?**

Multiple choice question with the following propositions:

- Undergraduate/Bachelor student
- Graduate/Master student
- Doctoral researcher
- Postdoctoral researcher
- Principal investigator/Group leader
- Assistant professor
- Associate professor
- Full professor
- Other (open field)

**19. Are you attending or have attended the ISRR11-ROOTING2021 conference?**

Close-ended question [Yes/No]

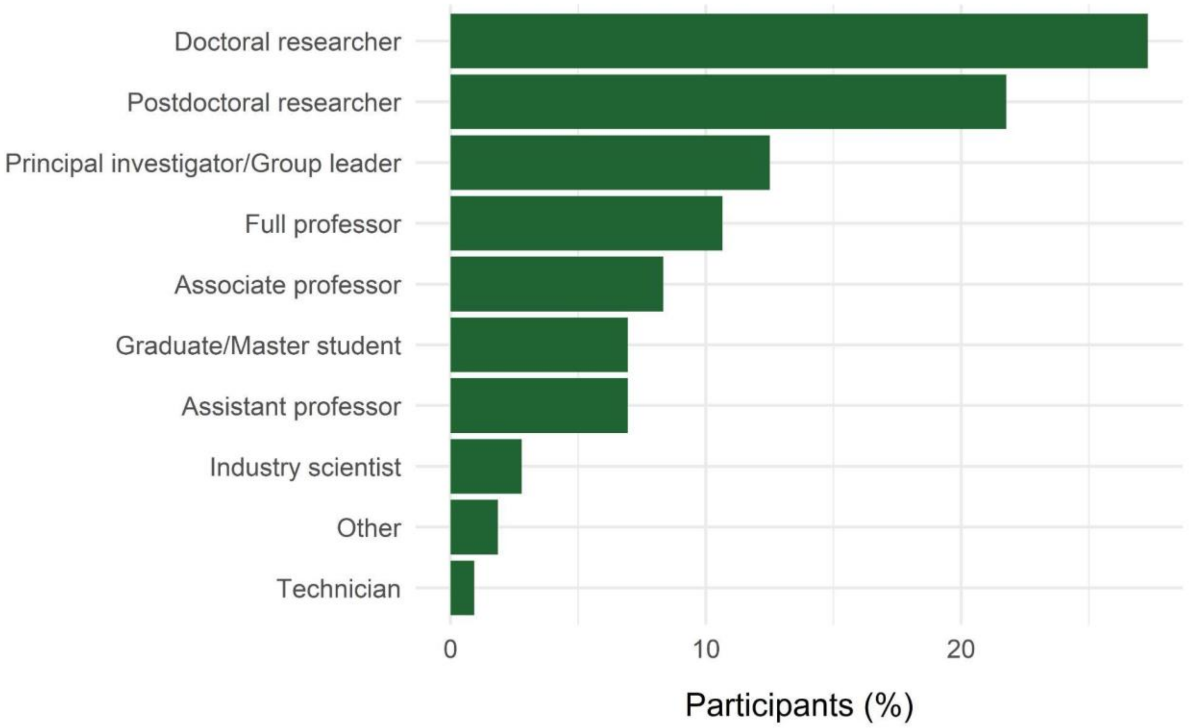


**Fig. S1**. Detailed description of the career stages of the participants in the online root phenotyping survey (*n*=216). The "Other" field includes career stage descriptions indicated by survey participants that could not be easily classified into one of the existing categories (e.g., early career research scientist, research scientist, senior researcher).

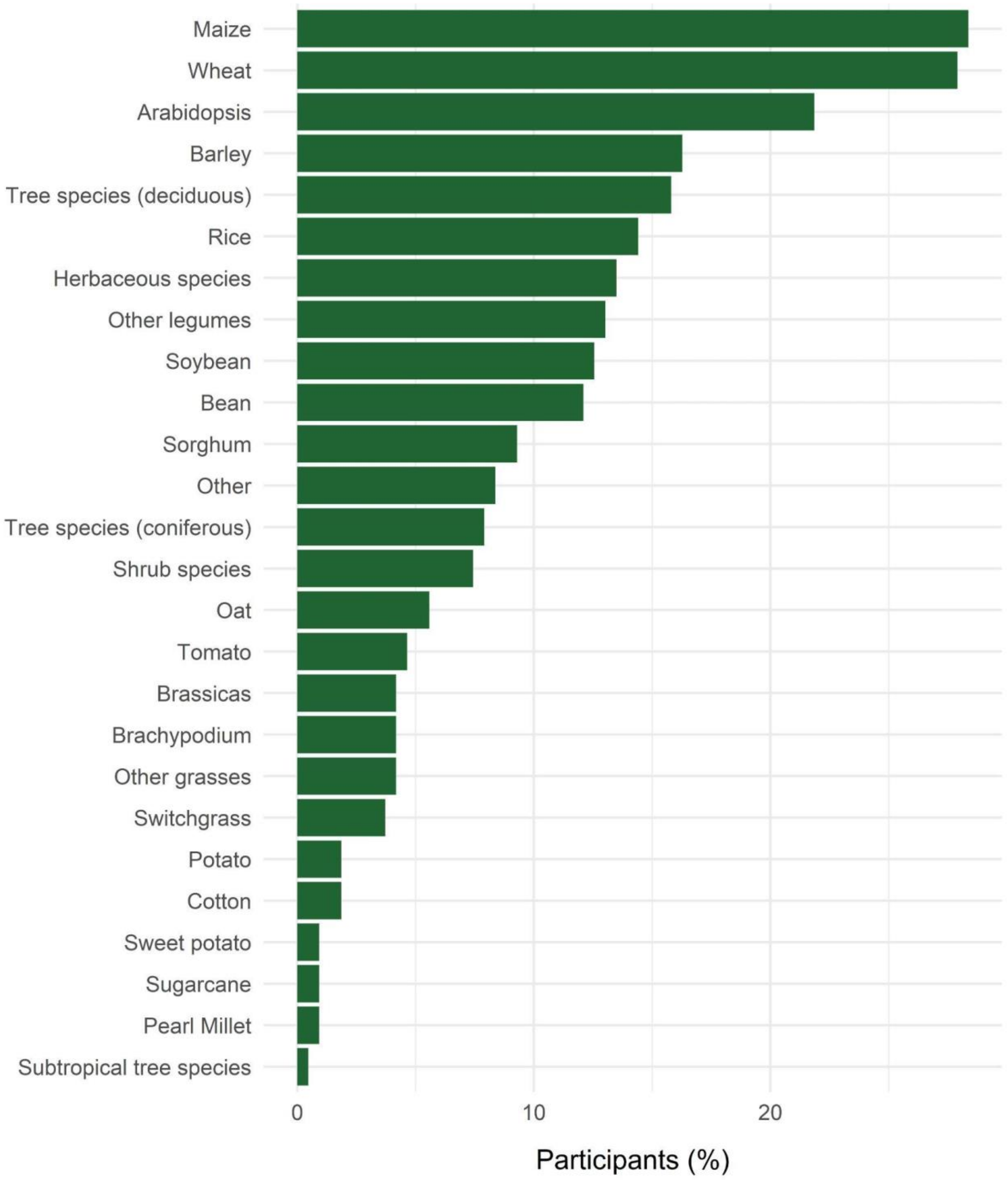


**Fig. S2**. Detailed description of the plant species most commonly used in root phenotyping studies according to the results of the online survey (*n*=215). For clarity, we grouped some of the species or groups of species provided by the survey participants into broader categories. The number of times a species or group of species was mentioned in the survey is indicated in brackets. The "Other legumes" category includes the following species or groups of species: cowpea (5), *Medicago truncatula* (3), chickpea (3), *Lotus japonicus* (2), clover (2), white lupin (1), white clover

(1), *Pisum sativum* (1), pigeon pea (1), persian clover (1), peanut (1), *Medicago* (1), lentil (1), (grain) legumes (4) and alfalfa (1). The “Other grasses” category includes the following species or groups of species: perennial ryegrass (2), pearl millet (2), timothy (1), *Setaria viridis* (1), ryegrass (1), orchard grass (1), finger millet (1), durum wheat (1) and bermudagrass (1). The “Brassicas” category includes the following species or groups of species: canola (3), Brassicas (2), watercress (1), pennycress (1), oilseed radish (1) and mustard (1). The “Other” category includes vegetables (1), tobacco (1), sunflower (1), *Plantago erecta* (1), phacelia (1), morning glory (1), linseed (1), *Lansium domesticum* (1), grapevine (1), cocoa (1), chicory (1), cassava (1), buckwheat (1), black pepper (1), basil (1), aubergine (1) and amaranth (1). Poplar (1), peach trees (1), *Citrus* trees (1) and apple trees (1) were included in the “Tree species (deciduous)” category.

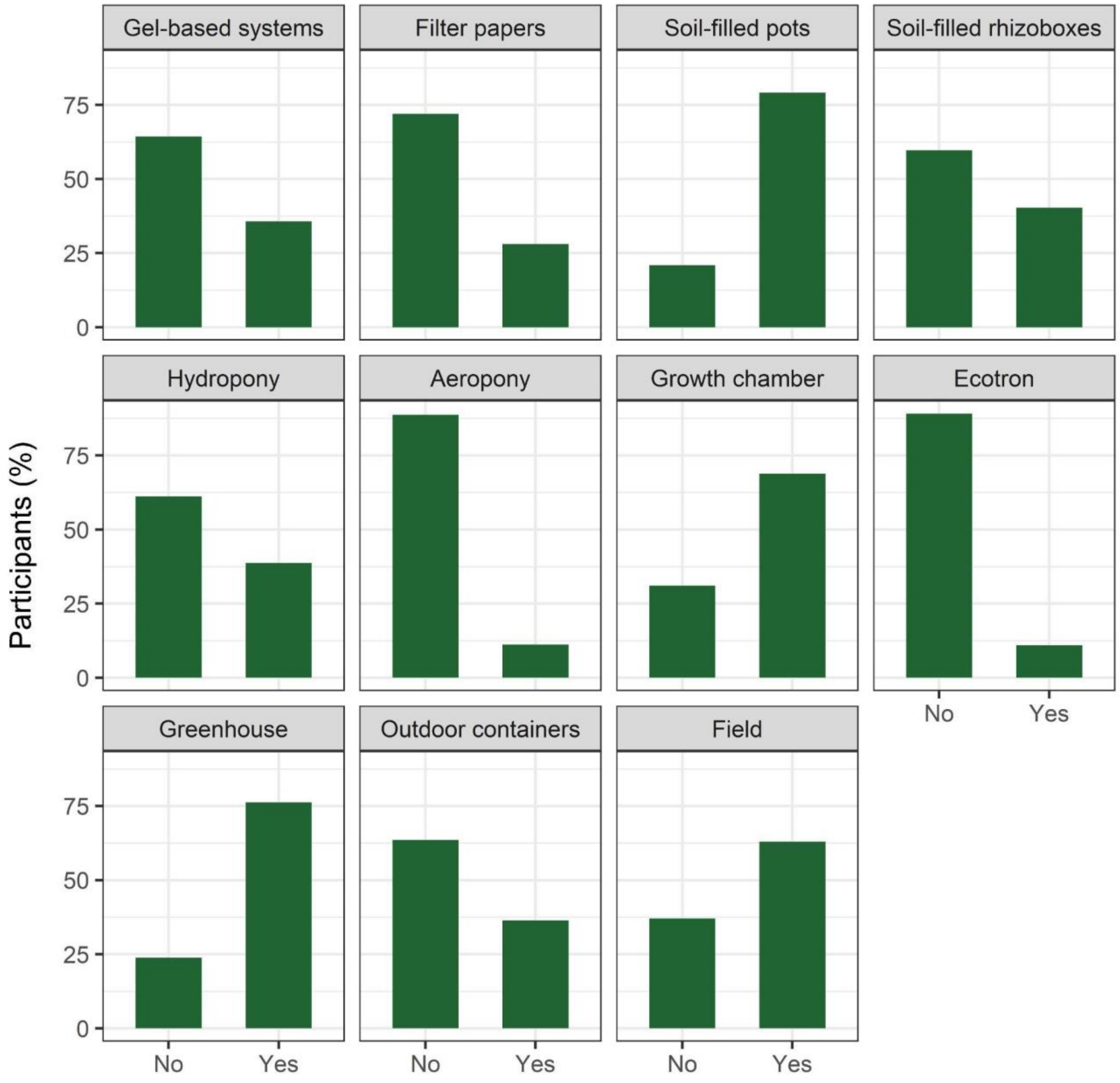


**Fig. S3**. Range of growing conditions used to grow plants in root phenotyping studies, according to the results of the online survey (*n*=216).

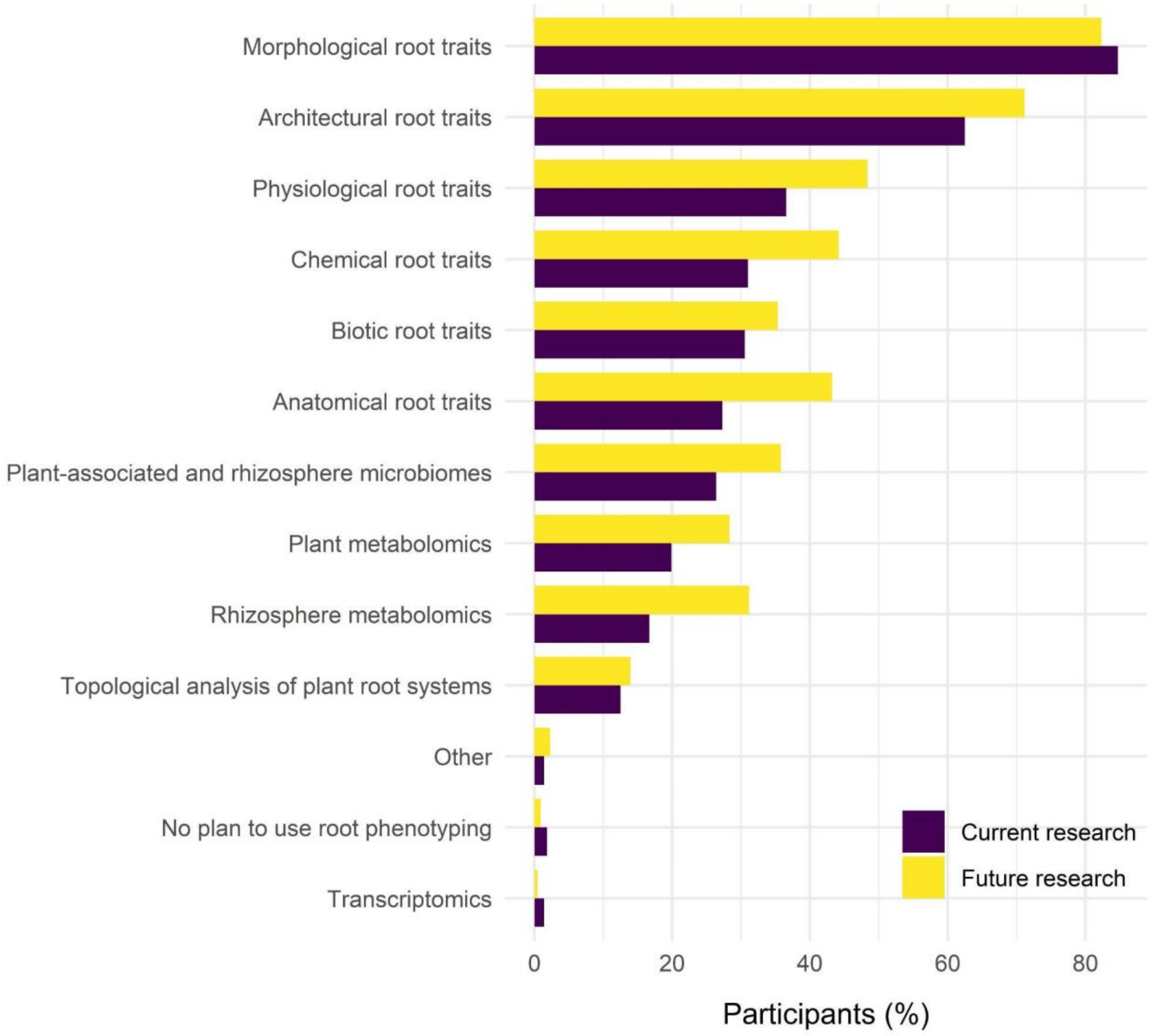


**Fig. S4**. Distribution of the main reasons why survey participants use or plan to use root phenotyping in their current and future research (current research: *n*=216; future research: *n*=215). The "Other" category includes reasons indicated by survey participants that could not be easily classified into one of the existing categories (acclimation to prevailing wind, plant ontology and metadata analysis, effects of fungicides on plant development, root-soil interface (macropores), root hair phenotyping).

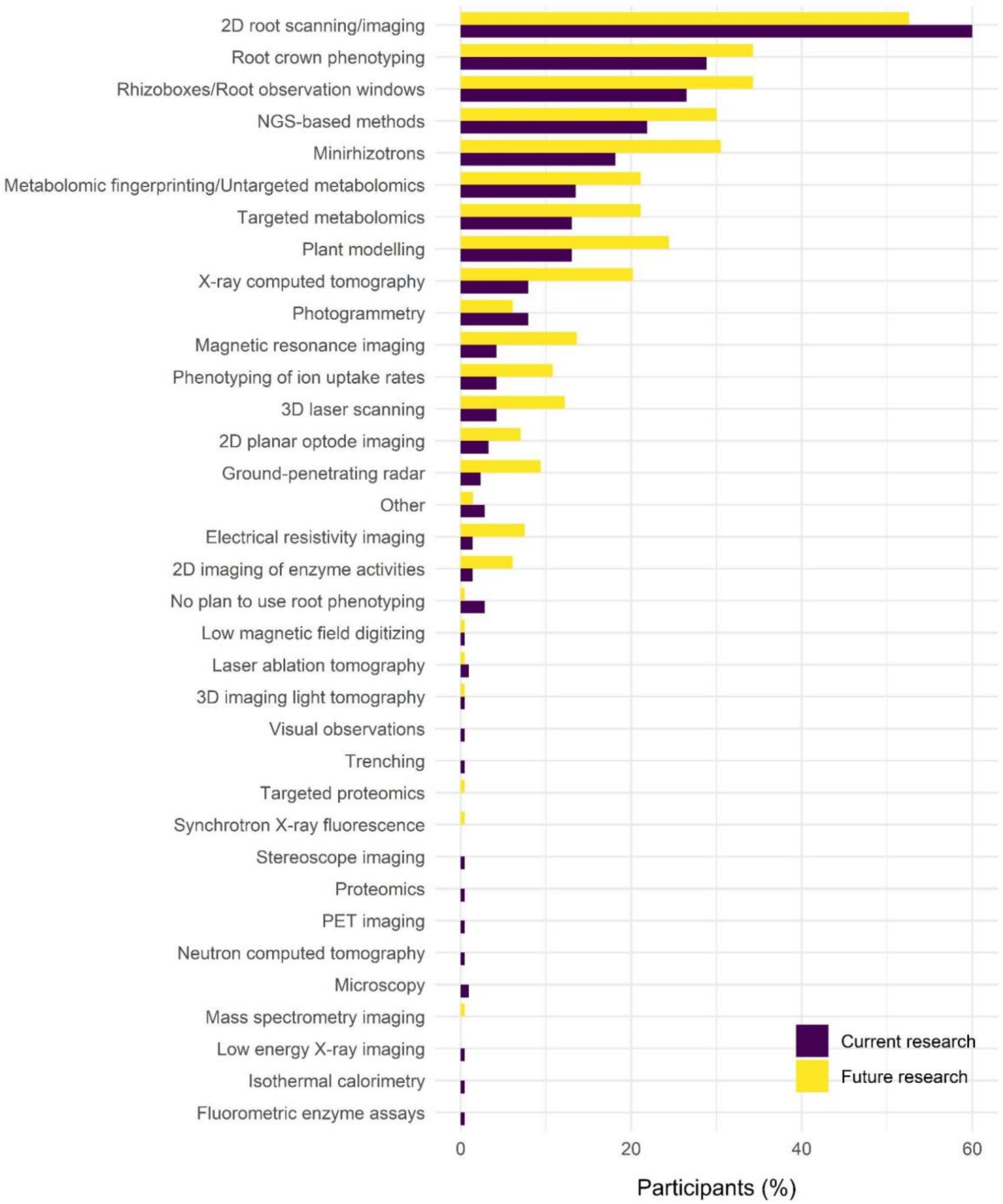


**Fig. S5**. Distribution of the main root phenotyping techniques that survey participants use or plan to use in their current and future research (current research: *n*=215; future research: *n*=213). The "Other" category includes techniques indicated by survey participants that could not be easily classified into one of the existing categories (plant ontology and metadata analysis, soil coring, dye-based macropore excavations, isotope based imaging).

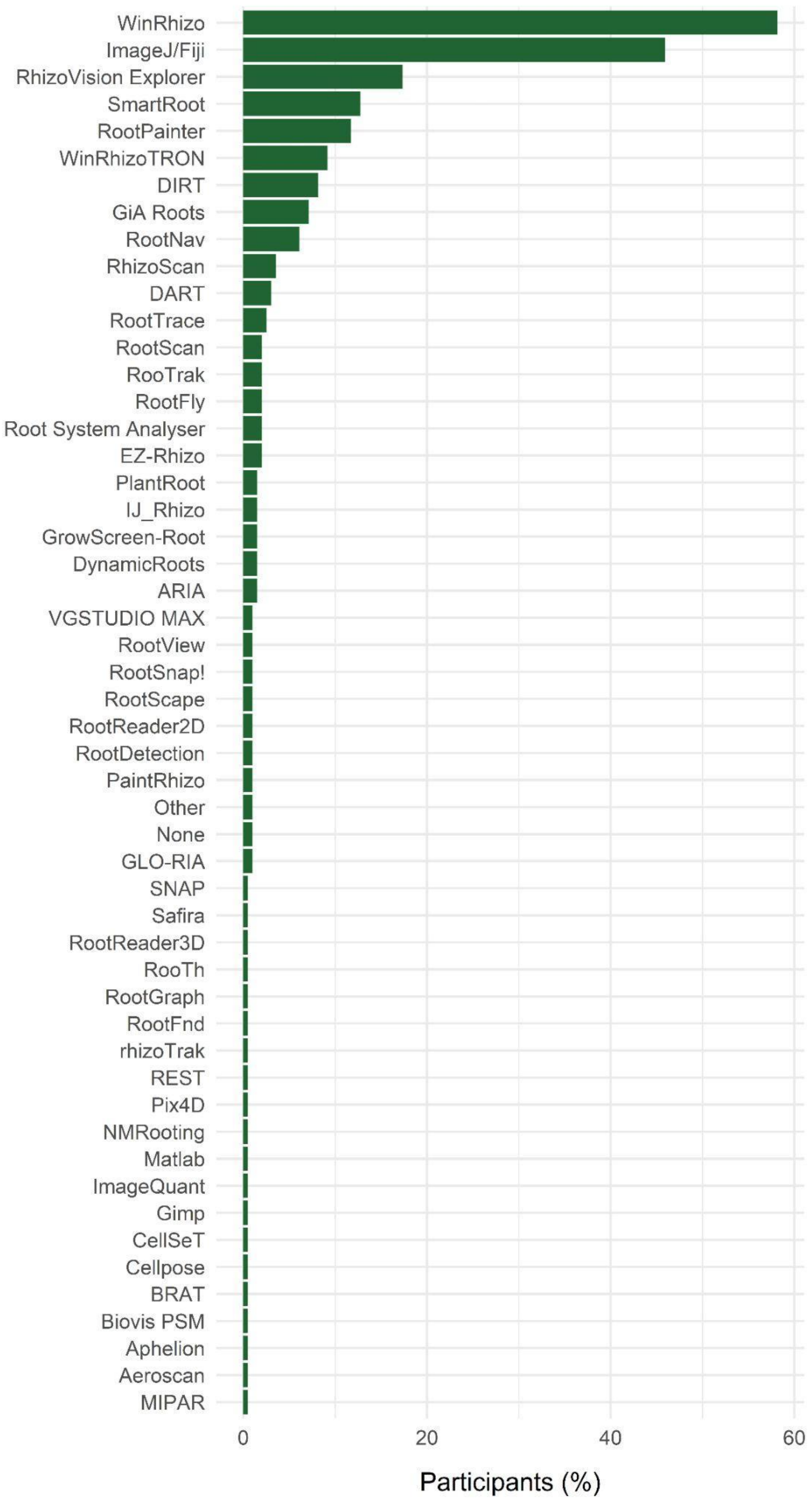


**Fig. S6**. Root image analysis software tools used in phenotyping studies (*n*=196).

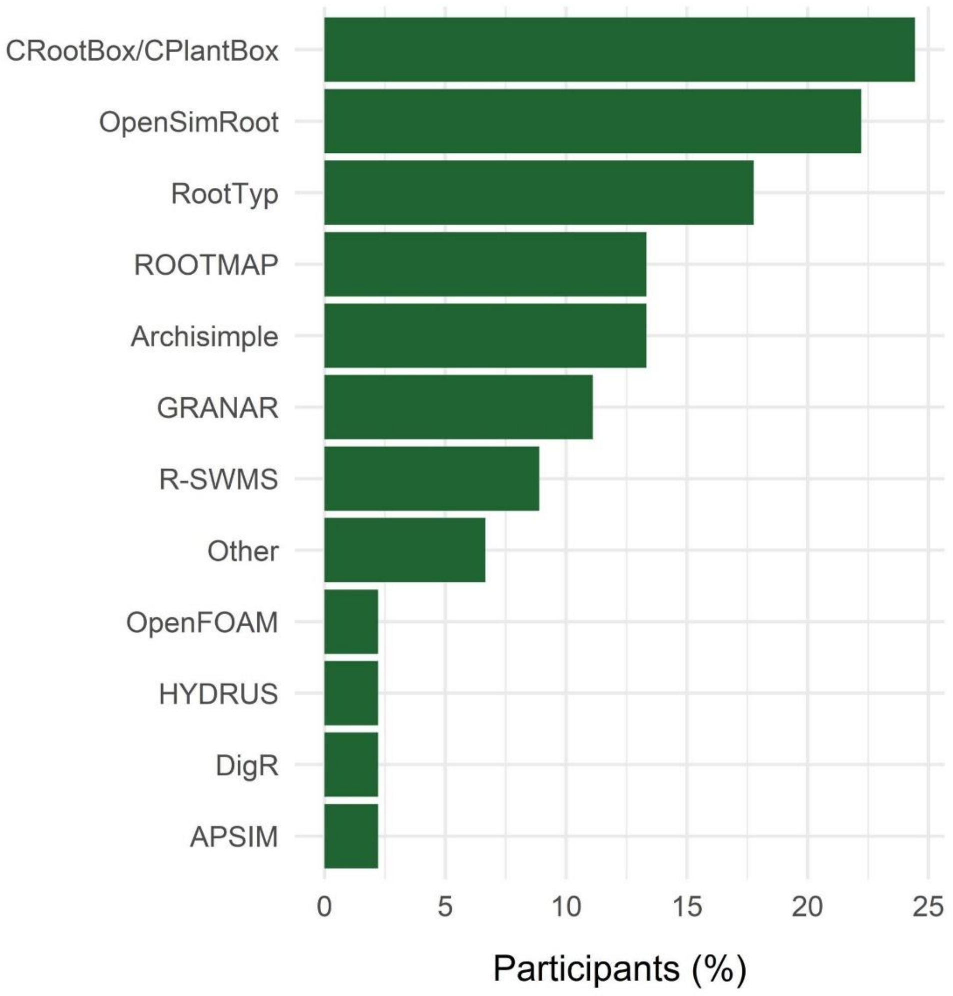


**Fig. S7**. Models used by survey participants in their plant modelling work (*n*=45). The “Other” category includes models indicated by survey participants that could not be easily classified into one of the existing categories (model proposed by Schenk and Jackson (2002), auxin distribution).

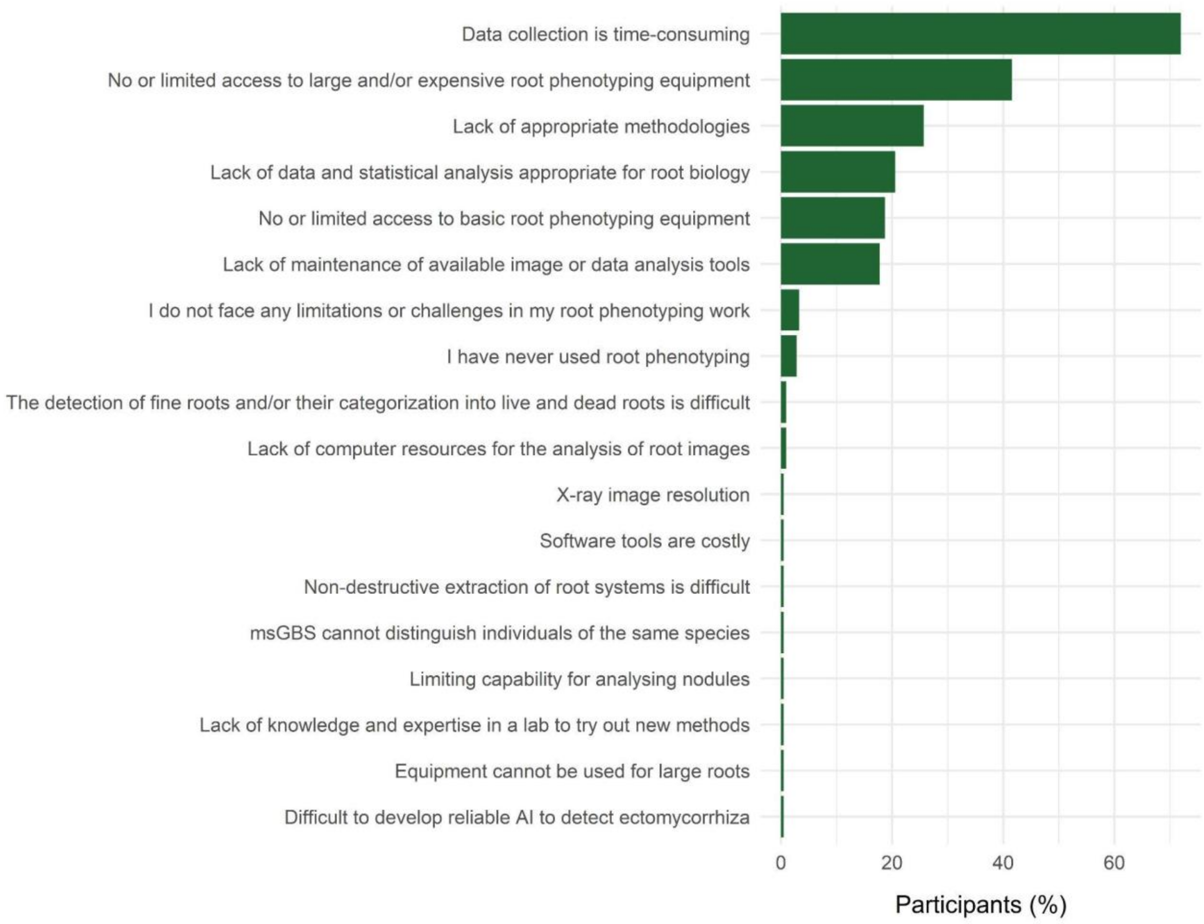


**Fig. S8**. Limitations and challenges encountered by survey participants in their root phenotyping work (*n*=214).

**Note S2.** The root scientists' wish list.

In this note, we attempt to summarise responses provided by participants of an online survey addressing the following question: What do you see as the most pressing developments in the field of root phenotyping that need to be made to enable you to carry out your future research plans? We added a few notes to redirect readers to useful information on some existing tools or some articles relevant to the issue in question.

***Sampling of roots, root exudates and associated microbes***

- Methods allowing a fast collection of soil and root samples in the field
- Methods allowing the non-destructive and 3D sampling of roots in the field
- Methods for the non-destructive sampling of rhizosphere and/or endosphere microbial communities
- Methods for the in situ collection of root exudates. ***Note***: See (Oburger & Jones, 2018; Williams *et al.*, 2021) for a detailed overview and comparison of root exudate collection techniques.
- Standardised protocols for the sampling of roots and root exudates in the field. ***Note***: See (Freschet *et al.*, 2021a) for a description of standardised measurement protocols in root ecology.

***Sample preparation***

- Fast, automated and efficient root washing techniques

***Phenotyping technologies, including image acquisition and analysis***

- Advanced but affordable imaging and analysis tools for in situ data collection (i.e., non-destructive, automated, high-throughput and accurate methods for phenotyping shallow and deep roots in the field and at all stages of plant development)
- New ways to monitor root growth non-destructively in systems that better represent the natural growing environment, including systems suitable for large mesocosms or tree species
- High-throughput methods for the measurement of root-associated functions. ***Note***: See (York, 2019) for a description of a functional phenomics pipeline. See also (Griffiths *et al.*, 2021) and (Guo *et al.*, 2021) for examples of platforms allowing the high-throughput phenotyping of ion uptake rates and root respiration, respectively.
- High-resolution methods for the in situ phenotyping of root hairs in different substrates
- Free, open-access, automated, accurate and easy to use image analysis software tools for root trait measurements (in 2D or 3D). ***Note***: See https://quantitative-plant.org/ for a

detailed list of root image analysis software tools meeting these criteria, including RhizoVision Explorer (Seethepalli *et al.*, 2021), RootPainter (Smith *et al.*, 2020; Han *et al.*, 2021) and DIRT (Das *et al.*, 2015).

- Root image analysis software tools that can be used on Windows, Mac and Linux computers.
- Fast and reliable methods to analyse (mini)rhizotron images, including the ability to track individual roots and quantify root turnover of different species based on morphological criteria such as root colour. ***Note***: See (Smith *et al.*, 2020) for a software tool allowing the segmentation of complex root images using deep learning.
- Imaging systems and image analysis software tools to track and analyse root development at different spatial resolutions (from anatomical features to entire root systems)
- Multi-plant imaging systems
- Methods for staining roots in vivo
- Small and affordable nutrient sensors
- Live cell imaging techniques that do not disrupt the structure of roots
- Higher resolution X-ray cameras and smaller spot size X-ray generators
- UAV (Unmanned Aerial Vehicle)-based root phenotyping methods in the field
- Methods allowing the quantification of root exudation rates in the field as well as methods to visualise metabolites in roots and soils
- Rhizosphere metabolomics pipelines allowing the identification and quantification of root exudates and soil metabolites. ***Note***: See (Uthe *et al.*, 2021) for a practical guide to implement metabolomics in ecological research.
- Image analysis-based methods for the quantification of mycorrhizal colonisation. ***Note***: See (Evangelisti *et al.*, 2021) for a software tool allowing the quantification of arbuscular mycorrhizal colonisation in plant roots using deep learning.
- Methods for the automatic counting of root nodules. ***Note***: See (Smith *et al.*, 2020) for a deep learning approach to count root nodules.
- New ways to track the growth of individual nodules and developing optical assays to monitor internal nodule processes
- Non-destructive methods to estimate the belowground biomass of large plant species such as trees
- New molecular techniques to distinguish individuals of the same species in mixed root samples
- Reducing the size of the equipment used in root phenotyping to make it easier to transport

and handle in the field

***Accessibility to root phenotyping equipment and facilities***

- Increasing the accessibility to basic root phenotyping equipment (e.g., flatbed scanners)
- Broadening the accessibility to high-throughput and/or expensive phenotyping and 3D imaging platforms such as X-ray computed tomography, magnetic resonance imaging (MRI), positron emission tomography (PET), and laser ablation tomography (LAT).
- Sharing knowledge on how to build phenotyping platforms

***Measuring root traits and functions***

- Standardised protocols for root trait measurements. ***Note***: See (Freschet *et al.*, 2021a) for a description of standardised measurement protocols in root ecology.
- Providing a mechanistic understanding of the roles of roots in plant and ecosystem functioning by exploring relationships between root traits (anatomy, physiology, morphology, architecture, biotic interactions) and functions (e.g., plant performance, nutrient and water uptake, resistance and tolerance to environmental stresses, etc.). ***Note***: See (Freschet *et al.*, 2021b) for a recent viewpoint on this topic.
- Improving species identification of roots in tropical forests based on morphological, physiological and biological descriptors
- Increasing the number of genotypes used in root phenotyping studies

***Modelling***

- Methods to help with the parameterisation of functional - structural plant models

***Data analysis***

- Data and statistical analysis tools appropriate for root biology
- Sustainable modelling and data analysis infrastructures, ontologies and formats. ***Note***: See (Lobet *et al.*, 2015) for a detailed description of the Root System Markup Language (RSML) file format.

***Data sharing***

- Sharing high quality, well-documented data to make it accessible to as many people as possible. ***Note***: Several databases allow researchers to share and reuse root trait data: FRED (Iversen *et al.*, 2017), TRY (Kattge *et al.*, 2011, 2020), GRooT (Guerrero-Ramírez *et al.*, 2021).

***Research funding and collaborative work***

- Funding programs to facilitate national and international collaborative networks
- Providing training for scientists who need root phenotyping in their research
